# Acoustic signatures in cavefish populations inhabiting different caves

**DOI:** 10.1101/2022.03.29.486255

**Authors:** Carole Hyacinthe, Joël Attia, Elisa Schutz, Didier Casane, Sylvie Rétaux

## Abstract

Complex patterns of acoustic communication exist throughout the animal kingdom, including underwater. The river-dwelling and the Pachón cave-adapted morphs of the fish *Astyanax mexicanus* share a repertoire of sounds, but their trigger, use and meaning has changed in cavefish after their recent colonization of the subterranean environment. Here, we explored whether and how sounds produced by blind cavefishes inhabiting different Mexican caves may have evolved, too. We compared acoustic parameters of sounds produced by wild cavefish, recorded in their natural settings, in six different caves located in three mountains ranges in North-East Mexico. Multi-parametric analyses show that the six cavefish populations sampled present cave-specific acoustic signatures, as well as possible individual signatures. The variations in acoustic parameters do not seem related to fish phenotypes, phylogeography or ecological conditions. We propose that the evolution of such acoustic signatures would be neutral and occur by drift, progressively leading to the differentiation of local accents that may prevent interbreeding and thus contribute to speciation.

## Introduction

Animal communication brings together all the information exchanges between individuals of the same or different species. The transmitter produces a signal, which causes a change in behavior or physiological state of the recipient. In the aquatic environment, where the speed of sound propagation is approximately four times faster than in the air and travels long distances, acoustic communication is widespread in mammalian and non-mammalian vertebrates such as teleosts [1–3]. In fishes, acoustic signals play roles in feeding, reproduction, hierarchy, predator detection, orientation and habitat selection.

Sonic animals have their own sound repertoires. The dolphin whistles, the whale clicks, or the toadfish boat-whistles are emblematic for aquatic species. Then, among species repertoires, individual signatures exist: Lusitanian toadfish males advertise their quality to females with boat-whistle calling rate [4], and both male and females frogs produce individual signature calls, suggesting that their acoustic communication may be more complex than expected [5]. Acoustic signatures are also species-specific. Their evolution has a suggested role in the speciation process, as proposed in cichlids [6, 7] or pipefishes [8]. In the later, differences in the structure of sound producing apparatus including cranial bone morphology may explain the unique acoustic signatures of the feeding clicks produced by closely related species. Also, within the piranha species *Serrasalmus marginatus*, red- and yellow-eyed morphs produce sounds with different amplitude features [9]. A mutation or different hormonal concentrations could explain both sound amplitude and eye color, playing a role in animal communication. This is, to our knowledge, a rare case of within-species acoustic signature in fish.

The fish *Astyanax mexicanus* (Mexican tetra) is also a sonic species [10]. Remarkably, acoustic communication has evolved between *Astyanax* river-dwelling and blind cave-adapted morphs, which have diverged about 20.000 years ago [11, 12]. Cavefish and river fish share a repertoire of six sounds, but functionally the trigger, the use, and the meaning of sounds has changed between cavefish from the Pachón cave and river fish [10]. Thus, acoustic communication has evolved after colonization of the subterranean habitat, opening avenues for the exploration of the evolution of acoustic signatures within the species.

## Results

In North-Eastern Mexico, there are more than 30 caves where cavefish populations leave [13, 14] and show signs of ongoing genetic differentiation [15–17], We performed acoustic recordings in March 2016 and 2017 in six different caves, in natural settings. We chose the Molino, Pachón, Los Sabinos, Tinaja, Chica, and Subterráneo caves because they are distributed throughout the 3 geographically distinct mountains ranges where *A. mexicanus* cavefish populations leave (**Fig. 1A**). Because of the topography and technical constraints encountered in each cave, the recording conditions, the number of fish recorded and the length of audio bands varied between caves [Molino (03/2017): 4 fish in openwork crate, 1h30; Pachón (03/2016): 10 fish in net, 11h; Los Sabinos (03/2017): 10 fish in net, 9h; Tinaja (03/2016), 25 fish in natural pool, 1h30; Chica (03/2017), 20 fish in natural pond, 11h; Subterráneo (03/2016): 12 fish in net, 10 hours; **Fig. 1B** and see Methods]. In agreement with known genetic or ecological conditions previously reported across caves e.g.[18], the phenotypes (size, level of troglomorphism) of the recorded fish were variable. For example, the fish recorded in Pachón cave were 2.7 to 6.7 cm in length (including one showing juvenile traits) and were fully troglomorphic, while the fish recorded in Subterráneo cave were 4.5 to 8 cm in length and among them two had tiny eyes (**Fig. 1C**).

**Figure 1:**
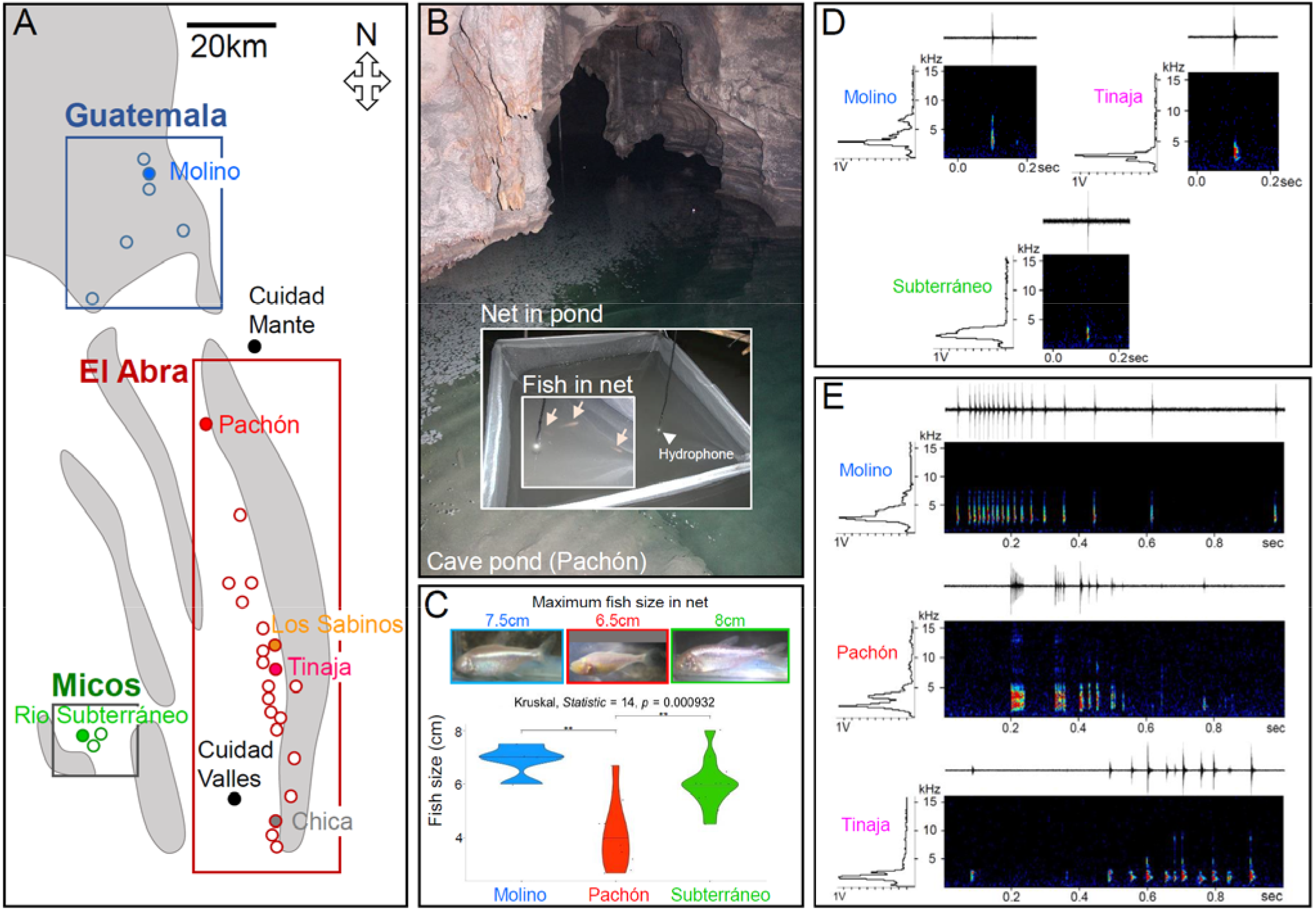
Sampling sounds in *A. mexicanus* caves. **A,** Map showing geographical localization of the 6 caves (color-coded) in 3 mountain ranges (rectangles) where acoustic production was recorded. **B,** Recording setup in natural settings. **C,** Comparison of the sizes of the fish recorded in different caves, with photograph of the largest individual present in the recording net. **D, E,** Examples of sonograms of Clicks (D) and Serial Clicks (E) recorded in the wild, showing substantial level of variations across caves.

A first, global analysis of audio bands (total 44h of recordings) showed that Clicks and Serial Clicks were the most represented sounds produced by cavefish in their natural environment, while the other sounds of the repertoire (Clocs, Serial Clocs, Sharp Clicks and Rumblings [10]) were rarer. We therefore focused on these two sounds, which are those showing the largest frequency bandwidth (500-10,000Hz) (**Fig. 1D,E**). We extracted and selected Clicks and Serial Clicks (n=50-100 each) for each cavefish population and compared their acoustic parameters (**Supplemental data, Table 1**).

Concerning single Clicks, we found that the sound duration, the dominant frequency, the signal to noise ratio/RMS power all varied significantly among the six caves (**Fig. 2A**). Clicks were longer in Tinaja (and Chica), more high-pitched in Molino (and Chica), and deeper and more powerful in Subterráneo. The signal to noise ratio was high in Chica and low in Tinaja. Of note, the duration and SNR variances were highest in Chica, suggesting less homogeneity in the sound production, possibly related to the hybrid genetic background of the fish in this cave. For Serial Clicks, we focused on the parameters related to their multi-pulse nature and the sound envelope. Pulse duration, number and rate, as well as interpulse and total sound duration were also variable among populations (**Fig. 2B**). Consistent with the single Click data, the pulse duration in Serial Clicks were longer in Tinaja. In Pachón, the pulse rate was impressively high, about ten times higher on average than in the other caves, accompanied by a number of pulses that was also twice higher, together with a shorter interpulse duration and a lesser total duration of the sound. Other significant features were high pulse numbers and long total sound durations in Subterráneo, and long interpulses in Molino. Together, these data show that sounds produced differ among caves, but with no apparent correlation or order relative to fish phenotypes (e.g., small Pachón fish can produce high pulse rates) or cave phylogeography (e.g., Pachón and Subterráneo fish share a high number of pulses in their Serial Clicks).

**Figure 2:**
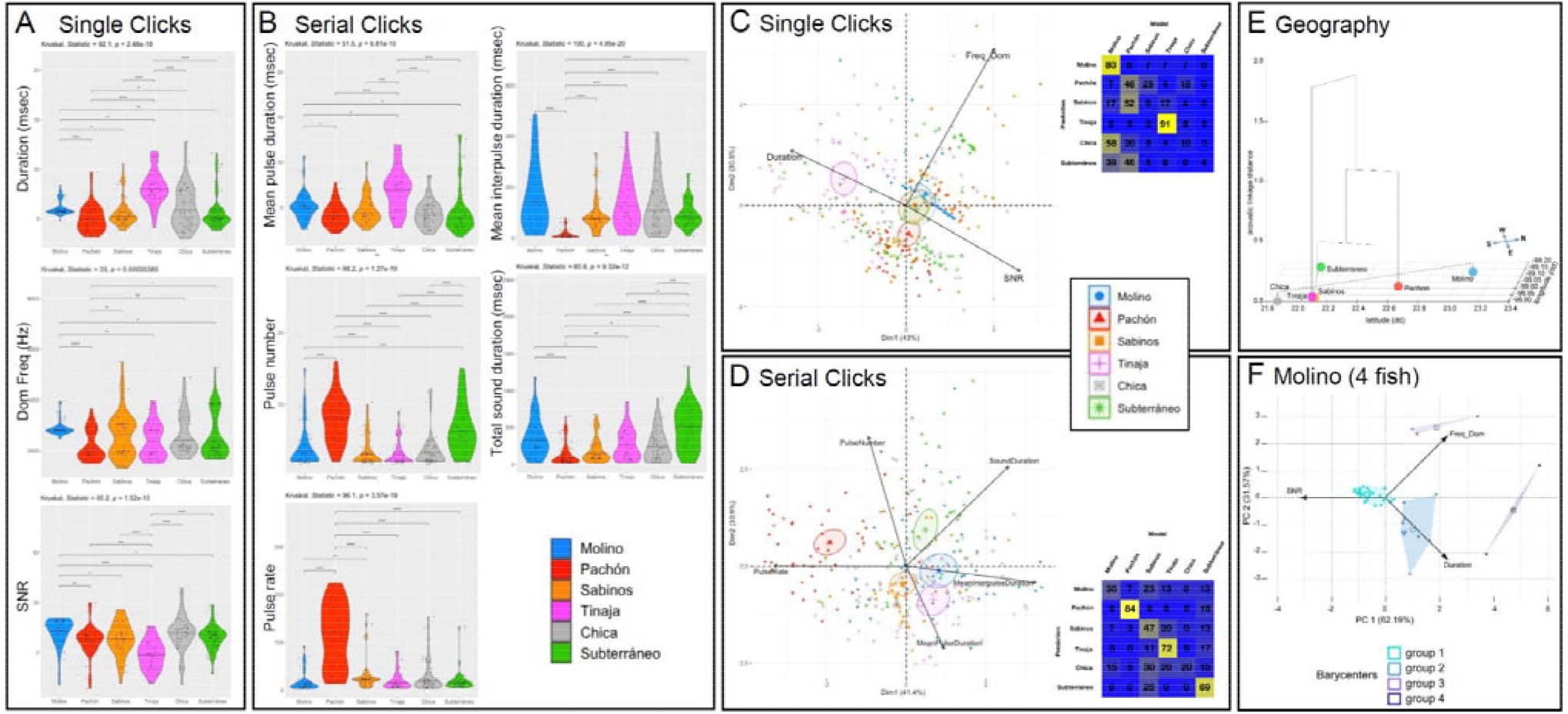
Comparing acoustic parameters of Single Clicks and Serial Clicks in *A. mexicanus* caves. **A-B,** Univariate analysis of acoustic parameters in different caves (color-coded). **C-D,** Multivariate analysis of acoustic parameters using PCA, followed by pDFA (insets). **E,** Hierarchical clustering of acoustic parameters of Single Clicks (height axis) projected onto a map of geographical coordinates of caves (NSWE in x and y-axes). **F,** Unsupervised clustering on PCA analysis for Single Clicks in the Molino cave (4 fish, 4 groups).

We next performed principal component analyses (PCA) to evaluate the possibility of a cave acoustic signature (**Fig. 2CD**). For both sounds, Clicks and Serial Clicks, confidences ellipses around centroids on the PCA showed little overlap (except Chica) and were mostly well separated. Moreover, pDFA (permutated Discriminant Function Analysis) generated confusion matrixes with good scores of correctly reclassified sounds, strongly suggesting that the sounds produced in each cave carry a specific acoustic signature **Fig. 2CD**, insets; p=0.001). Molino (80% reclassification), Pachón (46%) and Tinaja (91%) Clicks were particularly distinctive, and Serial Clicks from all caves except Chica showed significant reclassification scores (**Fig. 2CD**). Again, using a hierarchical clustering tree onto geographical coordinates of caves in the PCA (**Fig. 2E**, Single Clicks), we found some grouping of caves that does not fit with phylogeography. The apparent closeness of Subterráneo (Micos group) and Sabinos (El Abra group), or Molino (Guatemala group) and Chica (El Abra) rather suggests independent evolution of sound production in these different caves. In sum, our data provide strong evidence for the evolution of an “accent” in the different cavefish populations.

Finally, in our Molino dataset where only 4 fish were recorded, we noticed, using unsupervised hierarchical clustering on single Clicks, the presence of 4 different groups of sounds (**Fig. 2F**). Likewise, in Pachón (10 fish in net) and Subterráneo (12 fish in net) the same method yielded 8 groups and 9 groups of sounds, respectively (not shown). These results strongly suggest that besides their cave-specific acoustic signature, cavefish may also exhibit individual signatures.

## Discussion

The existence of acoustic signatures in recently evolved populations of cavefish was unexpected. Our findings provide an outstanding model to study the proximal and distal mechanisms for the evolution of acoustic communication in a species on its way to diversification and speciation [19] - even though *Astyanax* morphs still belong to the same species and show little genetic differentiation. The most likely origins for observed differences between caves include plasticity in response to specific local biotic and abiotic ecological conditions [13] see also [20], or the independent and subtle morphological evolution of facial and jaw bones in relation with the loss of eyes in cavefishes [21]. The latter would imply that the developmental evolution of the cavefish head not only affects the visual, olfactory and mechano-sensory but also the acoustic facet of their communication modalities, in a pleiotropic manner. We propose that the evolution of such acoustic signatures would be neutral and occur by drift, progressively leading to the differentiation of local accents that may ultimately prevent interbreeding and contribute to speciation.

In the subterranean environment too, the soft chirps of naked mole-rats encode individual identity as well as colony identity and they are culturally transmitted as colony vocal dialects, carrying information about group membership [22]. Although cavefish are supposed to be asocial, they are capable of social-like interactions in familiar environments [23]. The individual and cave-specific acoustic signatures, or accent, we have discovered may well participate in such sociality when thriving in their natural caves.

## Materials and Methods

### Fish samples

Field recordings were obtained during three field expeditions in the states of San Luis Potosi and Tamaulipas, Mexico, in March 2016 and March 2017, under the auspices of the field permit 02438/16 delivered by the Mexican Secretaria de Medio Ambiente y Recursos Naturales. We recorded from 6 caves hosting *Astyanax mexicanus* troglomorphic cavefish populations (map on Fig.1A).

### Sound recordings and analyses

For natural field recordings, cavefish were directly recorded in their hosting pools in 6 different natural caves, in the dark. This was done either from 10-12 fish maintained inside a large net (approx. 1m^3^ free water volume, Fig.1B) installed in their natural pool, or from freely swimming fish in the case of small natural pools (Tinaja and Chica caves). Hydrophones (Aquarian audio H2a XLR, Anacortes, WA, USA) were connected to portative pre-amplifiers (ART Dual Pre USB, NY 14305, USA) and recorders (Zoom H4n, NY 11788, USA) with SD cards, and recording parameters were adjusted with direct audio listening depending on environmental acoustic characteristics of each cave. They were left on sites for overnight recordings, except in the Molino cave: there, the entrance being a 70 meters vertical pit, the cave could not be visited on two consecutive days and the recording was limited to 1h30, during the afternoon. The four fish taped were nevertheless productive, as 50 clicks and 50 serial clicks could be extracted from this short period for analysis. Natural soundscape in Tinaja (heavy and continuous dripping limestone ceiling) prevented to exploit overnight recordings and made us seek for a quiet pond with specimens on the second day, leading to shorter recording as well (1h30). We therefore analyzed ≤100 sounds from the audio bands from other caves, to equilibrate the comparisons and statistical analyses with the more limited Molino and Tinaja dataset. Of note, for each cave the number of sounds analyzed (50) corresponded to minimum ten times the number of variables studies, i.e., 5 variables for the serial slicks.

Sounds were extracted by ear from audio bands recordings. Each audio track was carefully scrutinized manually from sonograms magnified at a 3-4 seconds temporal window and further at a 0.2 and 1 second bins (Fig.1D) allowing a visual control of each extracted sound motif while listening from T0 to end, including tracks of 11h. Clicks and serial clicks were not extracted in an exhaustive but in a systematical manner: only sounds previously described in the acoustic range of single click or serial click produced by fish in the lab or in the wild were considered, and replicates of each identified sound motif were randomly extracted. This allowed to increase the stringency and to capture the diversity of clicks and serial clicks for a comprehensive view of each cave soundscape.

Sounds were digitized at 44.1 kHz (16-bit resolution) and analysed using fast Fourrier transform (FFT) with Avisoft SAS Lab Pro 5.2.07 software (Avisoft bioacoustics, Glienicke, Germany) [24]. The acoustic structure of single clicks was analyzed using a set of:

-1 temporal parameter, the duration, measured from the envelop of the oscillogram,
-2 parameters measured from the oscillogram: RMS amplitude, and signal to noise ratio (SNR; [RMS amplitude of the signal-RMS amplitude of noise]/RMS amplitude of noise),
-5 spectral parameters obtained from power spectra (FFT, window type: Hann, window size: 512; time overlap: 90%) within a 0-22.5 kHz bandwidth. Spectral parameters were: 1) peak frequency of the frequency spectrum (dominant frequency), 2) first quartile of energy (Q25), i.e. the frequency value corresponding to 25% of the total energy spectrum, 3) second quartile of energy (Q50), 4) third quartile of energy (Q75), and 5) interquartile, i.e. difference between Q75 and Q25.

Serial clicks were examined using 5 fine temporal parameters including sound duration, pulse number, mean inter-pulse duration, mean pulse duration, and pulse rate (= sound duration/pulse number) using personal routines developed with R package Seewave [25].

Pulses were considered “single” if they were of short duration (<20msec) and separated by >1sec interval from the next pulse (threshold defined from the histogram of the inter-pulse durations). After calculating the correlation coefficients between the variables, we excluded highly correlated variables (r > 0.65 or < −0.65), retaining 3 uncorrelated variables for single clicks (duration, SNR, dominant frequency) and 5 uncorrelated variables for serial clicks. A principal component analysis (PCA; R package FactoMineR) was performed using the 3 variables retained for single clicks and the 5 variables retained for serial clicks, allowing to draw the centroïds of the 6 caves, surrounded by their 95% confidence circles (Fig.2CD). A permutated discriminant function analysis (pDFA; R routine from Bertucci et al. 2010 [24]) performed on the principal components axis of the previous PCA provided a classification procedure that assigned each sound to its appropriate cave (correct assignment) or to one of the others (incorrect assignment) (Fig. 2CD, insets).

Acoustic distances between caves plotted on a plan according to their actual GPS coordinates were calculated using an agglomerative hierarchical clustering on principle components (Euclidean metric, Ward method), for Single Clicks (Fig. 2E).

An unsupervised hierarchical clustering method was used on the Single Click Molino dataset and displayed on the ACP plan to estimate a potential individual acoustic signature in this cave (Fig.2F).

### Statistics

Normality was assessed with a Kolmogorov-Smirnov test. Kruskal-Wallis tests followed with Dunn’s *post hoc* were performed on non-normally distributed data sets. In figures, box plots show the distribution, median and extreme values (top and bottom whiskers) of samples. Statsoft Statistica 6, GraphPad Prism 9 and R 3.1.3 [26] were used for statistical analyses and graphical representations.

### A Source data file

(Supplemental data Table 1) containing all the raw data presented and analyzed in this paper is available as Supplemental Information.

## Acknowledgments

Work supported by a Lidex Neuro-Saclay collaborative grant to SR and JA, an Equipe FRM grant (DEQ20150331745) to SR, a prize from Fondation des Treilles to CH, and an Ecos-Nord exchange Program to SR and Patricia Ornelas-Garcia. We thank Luis Espinasa, Julien Fumey, and Stéphane Père and all other members of the Rétaux’s lab for their helpful spirit in the field, and Patricia Ornelas-Garcia for obtaining shared fieldwork permits.

